# Dimeric assembly of F_1_-like ATPase for the gliding motility of *Mycoplasma*

**DOI:** 10.1101/2024.06.11.597861

**Authors:** Takuma Toyonaga, Takayuki Kato, Akihiro Kawamoto, Tomoko Miyata, Keisuke Kawakami, Junso Fujita, Tasuku Hamaguchi, Keiichi Namba, Makoto Miyata

## Abstract

Rotary ATPases, including F_1_F_o_- and V/A-ATPases, are molecular motors that exhibit rotational movements for energy conversion^1^. In the gliding bacterium, *Mycoplasma mobile*, a dimeric F_1_-like ATPase forms a chain structure with the glycolytic enzyme, phosphoglycerate kinase (PGK), within the cell^2^, which is proposed to drive the bacterial gliding motility^2–4^. However, the mechanisms of force generation and transmission remain unclear. Here, we present a 3.2 Å resolution structure of the dimeric ATPase complex obtained using electron cryomicroscopy (cryo-EM). Notably, the structure revealed an assembly distinct from that of known dimeric forms of F_1_F_o_-ATPase^5^, despite containing conserved F_1_-ATPase structures. The two ATPase units were interconnected by GliD dimers, previously identified as MMOB1620^2,6^. Gliβ, a homologue of the F_1_-ATPase catalytic subunit^6^, exhibited a specific N-terminal region that incorporates PGK into the complex. Structural conformations of the catalytic subunits, catalytically important residues, and nucleotide-binding pattern of the catalytic sites of the ATPase displayed strong similarities to F_1_-ATPase, suggesting a rotation based on the rotary catalytic mechanism conserved in rotary ATPases^1,7–10^. Overall, the cryo-EM structure underscores an evolutionary connection in the rotary ATPases and provides insights into the mechanism through which F_1_-like ATPase drives bacterial gliding motility.

## Introduction

Rotary ATPases are molecular motors that link ATP hydrolysis or synthesis with ion transport via rotational motion in biological membranes^1^. These motors comprise two common domains: soluble R_1_, which is responsible for ATP hydrolysis and synthesis, and membrane-embedded R_o_, which is responsible for ion transport. F_1_F_o_-ATPase, one of the rotary ATPases conserved across most eukaryotes, archaea, and bacteria, is responsible for ATP synthesis and the maintenance of membrane potential^5^. An F_1_-like ATPase gene cluster, referred to as Type 2 ATPase, was reported in four *Mycoplasma* species, *M. mobile*, *M. pulmonis*, *M. agassizii*, and *M. testudineum*, in addition to the genuine F_1_F_o_-ATPase gene cluster, referred to as Type 1 ATPase^11,12^ (Extended Data Fig. 1). The presence of F_1_-ATPase α and β subunit homologues (MMOBs 1660 and 1670 in *M. mobile*) in the Type 2 ATPase gene cluster suggests a shared evolutionary origin in the F_1_ domain. Type 2 ATPase is proposed to drive the gliding motility observed in *M. mobile* and *M. pulmonis*^2–4,13,14^. *M. mobile*, a fish pathogen, glides at speeds of up to 4 µm per second on a solid surface covered with sialylated oligosaccharides found on host cell surfaces^15^ (Fig. 1a, Extended Data Fig. 2). In our proposed model for gliding, filamentous leg structures on the mycoplasma cell surface, which are approximately 50 nm, propel the cell body by catching and pulling sialylated oligosaccharides on the solid surface. Inside the cell, motors composed of Type 2 ATPase and phosphoglycerate kinase (PGK), which converts ADP to ATP through glycolysis, are linked near the cell membrane to form ‘internal chains’^2,6^. The motor is formed by dimerised F_1_-like ATPase units with unique bridge structures and is referred to as a twin motor. However, the mechanisms of force generation and transmission remain unclear. This study aimed to elucidate the detailed structure of the twin motor using cryo-EM single-particle analysis (SPA). Notably, the structural features had a strong homology to those of F_1_-ATPase^7,8^, suggesting a rotary catalytic mechanism analogous to that of rotary ATPase^1,9,10^ for the gliding motility of *M. mobile*.

**Fig. 1.**
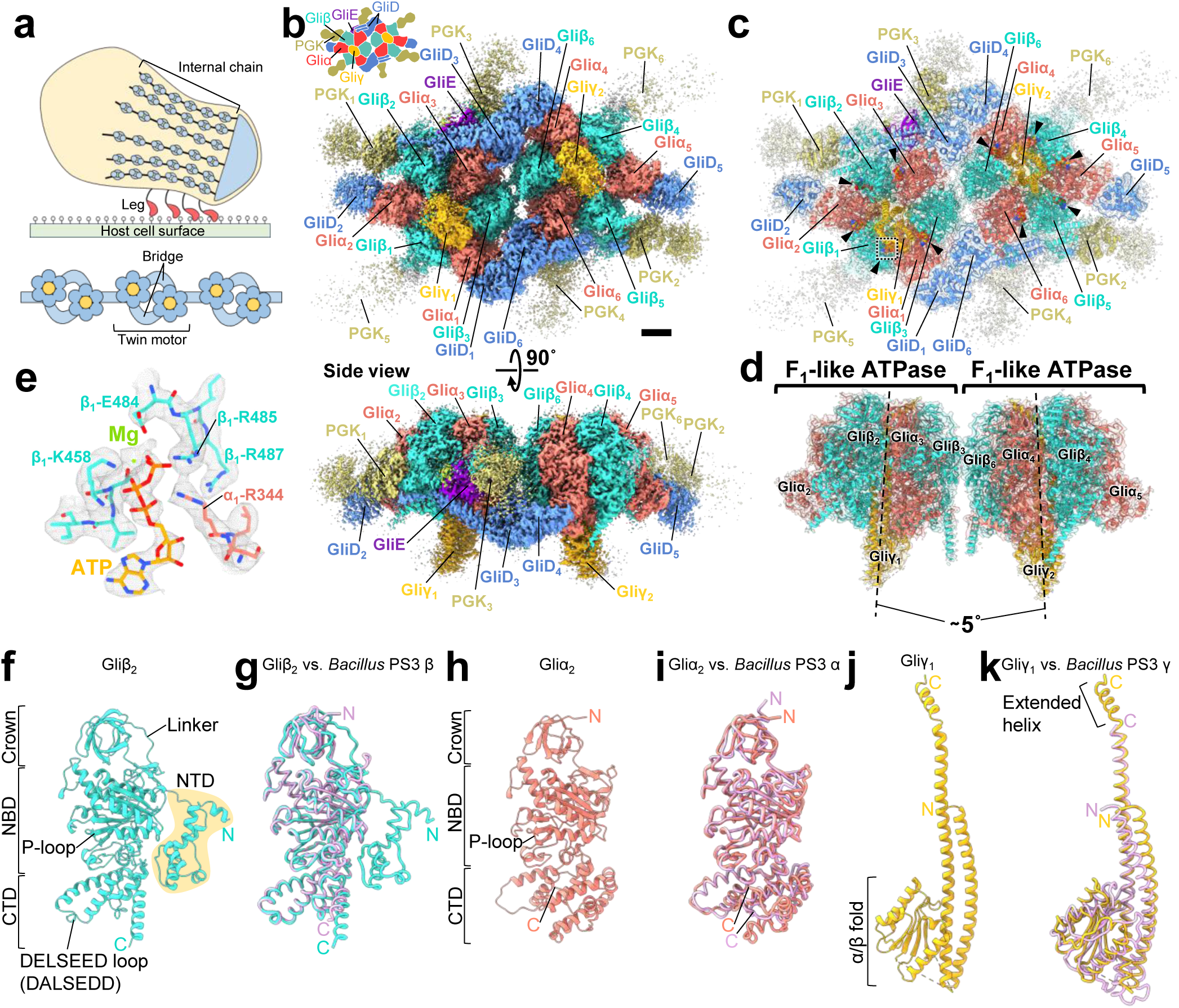
Structure of the twin motor featuring dimeric F_1_-like ATPase. (**a**) Gliding machinery of *Mycoplasma mobile*. Overall view of the gliding machinery (top) and part of the internal chain (bottom). (**b**) Overall structure of the twin motor. The illustration is drawn in the upper left. Scale bar = 25 Å. (**c**) Atomic model of the twin motor. Black triangles represent nucleotides. Maps in (b) and (c) are contoured at 0.55 in UCSF ChimeraX. (**d**) Dimeric F_1_-like ATPase in the twin motor. Dotted lines indicate the axis of each ATPase. (**e**) Density of Mg-ATP bound to F_1_-like ATPase. The dotted region in (c) is shown. (**f, h, j**) Subunits constituting F_1_-like ATPase. (**g, i, k**) Overlay of subunits of F_1_-like ATPase and *Bacillus* PS3 F_1_-ATPase. The subunits of the *Bacillus* PS3 F_1_-ATPase, αβ (PDB ID: 8HH5) and γ (PDB ID: 8HH2), are coloured purple.

## Results

### Protein identification using the cryo-EM structure of twin motor

Cryo-EM SPA was performed for twin motors isolated from *M. mobile* cells using images collected on the epoxidised graphene grid (EG-grid)^16^ (Extended Data Table 1). The overall 3D structure of the twin motor was determined at a resolution of 3.2 Å (Fig. 1b, Extended Data Figs. 3, 4). Local refinement improved the resolution of the regions, each containing the F_1_-like ATPase unit, to 3.1 Å (Extended Data Figs. 3b, 4). The density maps enabled the construction of the atomic model of the twin motor (Fig. 1c). The modelled sequence regions of each subunit are summarised in Extended Data Table 2. The twin motor exhibited a rectangular structure with dimensions of approximately 350 × 250 × 150 Å and at least 24 polypeptide chains with a total mass of approximately 1.5 MDa. Notably, the two F_1_-like ATPase units were inclined to each other by approximately 5° and formed a dimer with pseudo-two-fold rotational symmetry and several accessories (Fig. 1c, d). The F_1_-like ATPase comprised a hexameric ring of two alternating subunits and a central shaft subunit, identified as MMOBs 1660, 1670, and 1630, respectively. In the present study, these subunits were named Gliα, Gliβ, and Gliγ, according to the subunit names of F_1_-ATPase^10^. In the hexameric ring, nucleotide-derived densities were observed at five of the six subunit interfaces, despite the isolation and analysis of samples under nucleotide free condition, indicating the presence of endogenous nucleotides as reported in F_1_-ATPase^17^ (Fig. 1c, e). The subunit attached to Gliα, forming a ‘bridge’ between two F_1_-like ATPase units, was identified as MMOB1620 and named GliD in the present study (Fig. 1c). The three densities protruding from the dimeric F_1_-like ATPase were identified as MMOB4530, which was previously annotated as PGK. The denoised 3D map revealed additional distinct densities at six locations, including the PGK_1–3_ regions (Extended Data Fig. 5a). These six densities exhibited correlation coefficients ranging from 0.37 to 0.48 with the crystal structure of PGK from *Staphylococcus aureus* (PDB ID: 4DG5) (Extended Data Fig. 5b, c). Although the values are low, these assignments are consistent with the stoichiometry previously estimated via SDS-PAGE^2^, suggesting these maps are PGK molecules. A density bound only to one of the two F_1_-like ATPase units was identified as MMOB3660, which was newly detected via SDS-PAGE of the twin motor and named GliE in the present study (Fig. 1b, c, Extended Data Fig. 6). This small protein (112 aa), characterised by a β-sandwich, was nestled between the ATPase hexamer and other subunits, breaking the rotational symmetry of the complex (Extended Data Fig. 7). Although MMOB1640, encoded in the Type 2 ATPase gene cluster, was detected via SDS-PAGE of the twin motor (Extended Data Fig. 6), the protein density was not observed from the overall 3D map.

### Structural features of subunits comprising F_1_-like ATPase

Gliβ and Gliα exhibited high structural similarity to the β and α subunits of the F_1_-ATPase from the thermophilic bacterium, *Bacillus* PS3^8^ (PDB ID: 8HH5), based on the superimposition of their Cα atoms with the root mean squared deviation (RMSD) values of 1.95 Å and 2.79 Å, respectively (Fig. 1g, i). Both Gliβ and Gliα had three domains: crown, nucleotide-binding domain (NBD), and C-terminal domain (CTD). The NBDs of Gliβ and Gliα comprise a phosphate-binding motif of P-loop^18^, consisting of the amino acid sequences GGAGVGKT and GDRGTGKT, respectively (Fig. 1f, h). The Gliβ-CTD contained a DELSEED loop, which is important for torque transmission in the F_1_-ATPase β subunit^19^, with an amino acid sequence of DALSEDD (Fig. 1f). A remarkable difference between Gliβ and the F_1_-ATPase β subunit was the presence of an extended N-terminal region consisting of the N-terminal domain (NTD) and linker (Fig. 1f, Extended Data Fig. 8a). This region comprised approximately 100 residues located within the extra N-terminal 299 residues of Gliβ^6,12^. Notably, the linker hangs outside the F_1_-like ATPase along the crown and NBD (Extended Data Fig. 8b). Gliβ-NTDs, consisting of 4 to 6 helices, were oriented in various directions among Gliβ, depending on their position in the complex (Extended Data Fig. 8a, c, d). Further, Gliβ-NTD interacted with PGK (Fig. 2a, b). In addition, at the interface of the two F_1_-like ATPase units, the N-terminal regions of Gliβ_3_ and Gliβ_6_ interacted with the NBDs of Gliα_6_ and Gliα_3_ of another F_1_-like ATPase unit, respectively (Fig. 2a, c). These interactions suggest that the Gliβ-specific N-terminal region plays a role in the incorporation of PGK into the twin motor and stabilisation of the ATPase dimer. A difference between Gliα and the F_1_-ATPase α subunit is the absence of the H1 helix, which interacts with the peripheral stalk^20^ (Extended Data Fig. 9). This absence is consistent with the lack of a gene homologous to the peripheral stalk in the Type 2 ATPase gene cluster^12^. Gliγ, which is composed of a coiled coil and an α/β fold typical of the F_1_-ATPase γ subunit, was superimposed on the γ subunit of the *Bacillus* PS3 F_1_-ATPase with an RMSD value of 5.79 Å for Cα atoms due to a lower sequence conservation than that of the catalytic subunits^15^ (Fig. 1j, k). The folded C-terminal α-helix, which is 25 Å longer than the F_1_-ATPase γ subunit, protrudes from the N-terminal side of the hexameric ring and interacts with the crowns of Gliαβ (Fig. 1k, Extended Data Fig. 10a, b). The coiled coil of Gliγ hydrophobically interacted with the CTDs of Gliαβ in the hexameric ring (Extended Data Fig. 10c), and in the F_1_-ATPase, this interaction is involved in axle hold and torque transmission^21^.

**Fig. 2.**
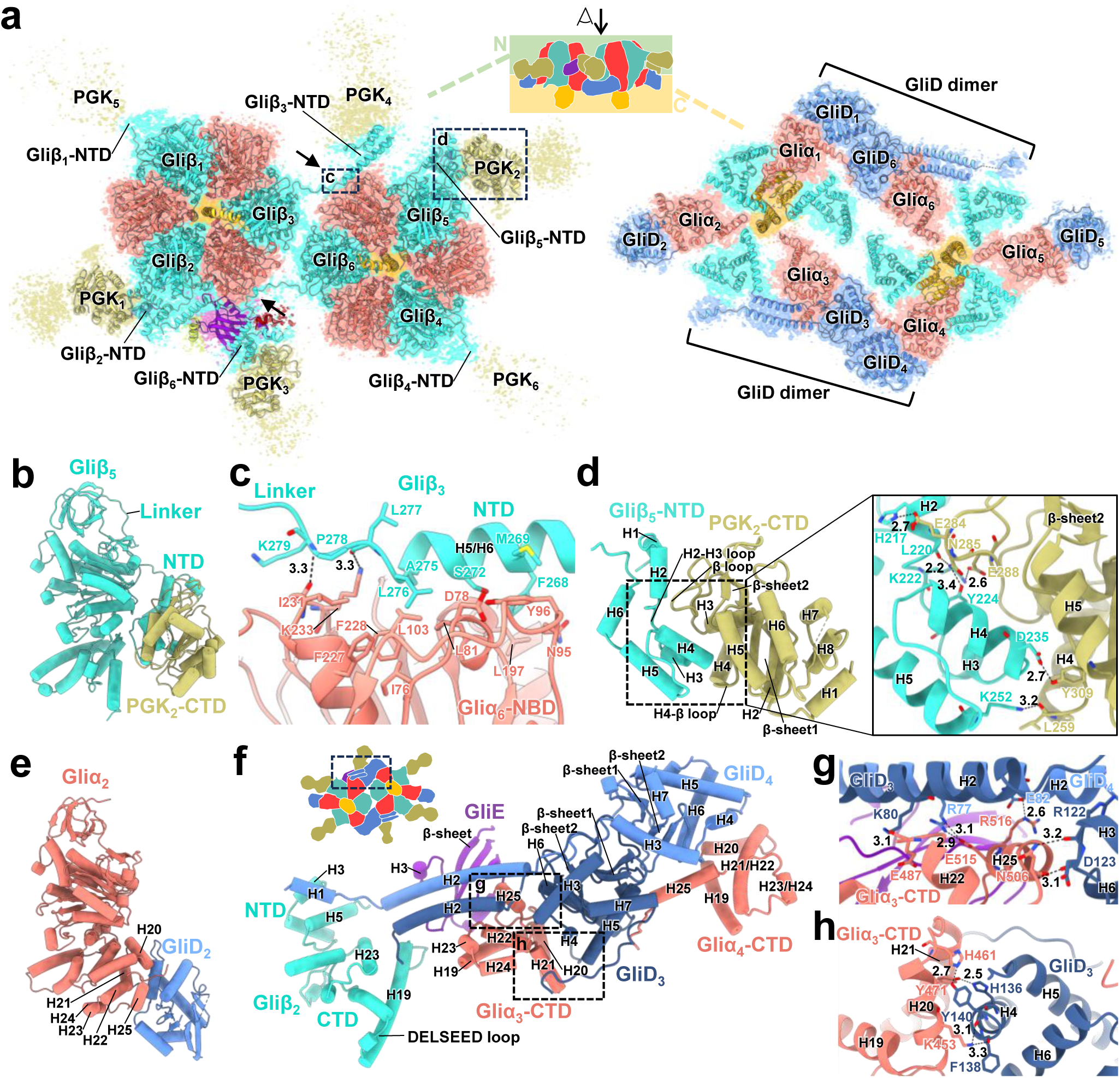
Interaction between subunits. (**a**) N- and C-terminal regions of the twin motor. The twin motor is viewed from the top of the illustration. Arrows indicate the interface between dimeric F_1_-like ATPases. Maps are contoured at 0.55 in UCSF ChimeraX. (**b**) Structure of Gliβ and PGK. (**c**) Interaction between the N-terminal region of Gliβ and Gliα-NBD at the interface between dimeric F_1_-like ATPases. The dotted region in (a) is shown. (**d**) Interaction of PGK with the Gliβ-NTD. The dotted region in (a) is shown. (**e**) Structure of the Gliα and GliD monomers. (**f**) GliD dimer at the interface between dimeric F_1_-like ATPases. The figure corresponds to the dotted region in the illustration. (**g, h**) Interaction between GliD and Gliα-CTD. Each corresponds to the dotted regions in (f).

### Accessories of dimeric F_1_-like ATPase

Generally, PGK is bifurcated into two domains interconnected by an α-helix: the NTD, which binds to 3-phosphoglycerate (3-PG) or 1,3-bisphosphoglycerate (1,3-BPG), and the CTD, which binds to Mg-ADP or Mg-ATP^22^ (Extended Data Fig. 11a). The twin-motor PGK structure, comprising the CTD and a segment of the α-helix, exhibited a high structural similarity to the PGK from *S. aureus* (PDB ID: 4DG5), with an RMSD value of 3.12 Å for Cα atoms (Extended Data Fig. 11b, c). The β loop of the PGK-CTD invades the depression of the folded Gliβ-NTD and interacts with the H2 helix and the H2–H3 loop of the Gliβ-NTD (Fig. 2d). The H4–β loop and H5 helix of the PGK-CTD also interacted with the H5 and H4 helices of the Gliβ-NTD, respectively.

Nucleotide-derived densities were not found in the PGK-CTD of the twin motor, despite conservation of the residues important for nucleotide binding in PGK^23^ (Extended Data Figs. 11d–f, 12). None of the twin-motor PGK-NTDs were modelled due to the poor densities of the reconstruction. However, the binding and catalytic residues for 3-PG or 1,3-BPG in PGK were conserved in terms of amino acid sequence, suggesting that the twin-motor PGK possesses enzymatic activity (Extended Data Fig. 12). As the binding and catalytic residues were not conserved in another PGK homologue, MMOB4490, encoded in the *M. mobile* genome, the twin-motor PGK is expected to function as the original PGK in glycolysis (Extended Data Fig. 12). The GliD (GliD_1,3,4,6_) bridge components formed dimers at the interface of two F_1_-like ATPase units, whereas GliD_2,5_at the twin-motor end existed as monomers (Fig. 2a, e, f). GliD contained a globular domain and an N-terminal long α-helix (H2), which was only visualised in the dimer. The globular domain interacted with the H20–H25 region at the C-terminus of Gliα (Fig. 2g, h). The two H2 helices in the GliD dimer aligned parallel to each other across the H20–H25 region of Gliα, GliE, Gliβ-NTD, and Gliβ-CTD (Fig. 2f). These two α-helices are reminiscent of the dynein buttress that connects the AAA+ ring and the tubulin-binding domain^24^, suggesting a regulatory role for structural changes induced by the ATPase activity of the F_1_-like ATPase units.

### Structural information suggesting a catalytic mechanism analogous to that of F_1_-ATPase

The crown region of the hexameric ring exhibited approximately six-fold rotational symmetry, whereas the CTD region was asymmetric (Extended Data Fig. 10a). This observation suggests a structural change with a pivot point between the crown and CTD. Gliβ_1–3_ adopted three distinct conformations, which appear to be the transition states of CTD flexion motion towards Gliγ, with a positional difference of 11 Å for the DELSEED loop (Fig. 3a). These conformations were named Gliβ_C_ (closed), Gliβ_HO_ (half-open), and Gliβ_O_ (open), according to the conformation names of F_1_-ATPase^8^. The conformations were arranged clockwise when viewed from the C-terminus of the hexameric ring (Fig. 3b). These conformations resemble the states referred to as *ATP-waiting* or *step-waiting*, which occurs during the catalytic reaction of *Bacillus* PS3 F_1_-ATPase; the latter occurs after ATP binding^8^ (Extended Data Table 3). In Gliα, the region corresponding to the DELSEED loop exhibited only a 5 Å difference in position relative to the Gliγ side, whereas the H20–H25 region interacting with GliD exhibited a difference of 9 Å (Extended Data Fig. 13). These conformations were named Gliα_C_ (close), Gliα_HO_ (half-open), and Gliα_O_ (open) based on the H20–H25 positions viewed from the Gliγ side. If this difference in Gliα represents a conformational change associated with ATP hydrolysis of the twin motor, force transfer may occur from F_1_-like ATPase to GliD. The catalytic sites formed at the α–β interface of Gliα_C_β_C_ and Gliα_HO_β_HO_ contained Mg-ATP and Mg-ADP + P_i_, respectively, whereas no nucleotide density was found at the catalytic site of Gliα_O_β_O_ (Fig. 3b–d, Extended Data Fig. 14).

**Fig. 3.**
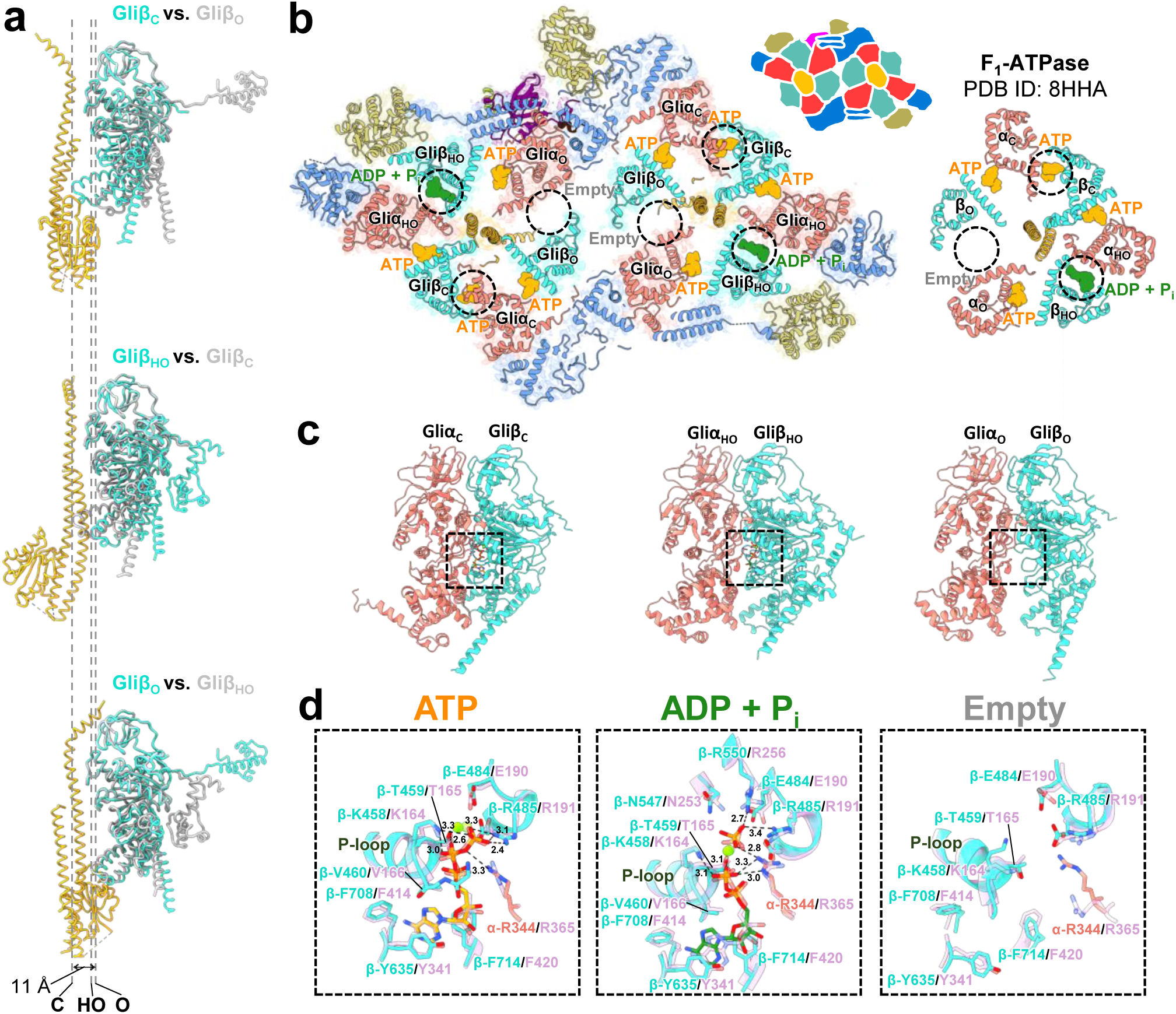
Structural conformations and nucleotide binding of Gliβ. (**a**) Differences in the conformations of the three Gliβ. Superposed three Gliβs in the F_1_-like ATPase. The dotted lines indicate the position of the DELSEED loop of each Gliβ. (**b**) Nucleotide binding pattern of dimeric F_1_-like ATPase and F_1_-ATPase. The catalytic site is circled by black dotted lines. (**c**) Gliαβ dimer in each conformation in the hexamer. (**d**) Comparison of the catalytic sites. Each catalytic site corresponds to the dot square above. The catalytic sites are superimposed on the corresponding catalytic sites of F_1_-ATPase (PDB ID: 8HHA), which are coloured purple.

The nucleotide-binding states were also shared between the two F_1_-like ATPases according to the pseudo-two-fold rotational symmetry. The residues of F_1_-ATPase involved in ATP hydrolysis, including those in the arginine finger, were conserved in Gliαβ, suggesting a shared ATP hydrolysis pathway between F_1_-ATPase and the twin motor (Fig. 3d). Mg-ATP was observed at the remaining three interfaces, hereafter referred to as the β–α interfaces (Fig. 3b, Extended Data Fig. 14). At the β–α interface, Gliα-K179 corresponded to the position of conserved residue Q200 of the F_1_-ATPase α subunit (Extended Data Fig. 15a, b). Compared with the α–β interface, Gliα-K179 is charge inverted relative to the glutamate essential for the catalytic activity (E190 of the F_1_-ATPase β subunit), suggesting that the β–α interface, akin to F_1_-ATPase, does not exhibit ATP hydrolysis activity (Extended Data Fig. 15c). The conformational sequence pattern of three Gliβ subunits with bound nucleotides was identical to that of the *ATP-waiting* state of F_1_-ATPase^8^ (Fig. 3b). These data suggest that Mg-ATP binding to the catalytic site of Gliα_O_β_O_ promotes the catalytic reaction of the entire F_1_-like ATPase based on the catalytic mechanism analogous to that of F_1_-ATPase.

### Twin motors in the chain

The atomic model of the twin motor was fitted into the previously reconstructed density of the internal chain, obtained using negative staining electron microscopy (EM) SPA^2^ (Fig. 4a). To circumvent misfitting of the long-axis orientation of the twin motors due to the low-resolution density map (29.7 Å) of the chain structure, regions with asymmetric structures, such as the NTDs of Gliβ_3,6_, GliE, and PGK_3_, were excluded from the fitted model. At the interface between twin-motor units, GliDs that project along the long axis of the twin motor faced each other, suggesting their role as a linker for chain formation. However, the 3D map of the internal chain revealed a remaining density at the interface that cannot be filled by GliDs, which may correspond to the approximately 100 residues comprising an N-terminal region and loops that have not been modelled.

**Fig. 4.**
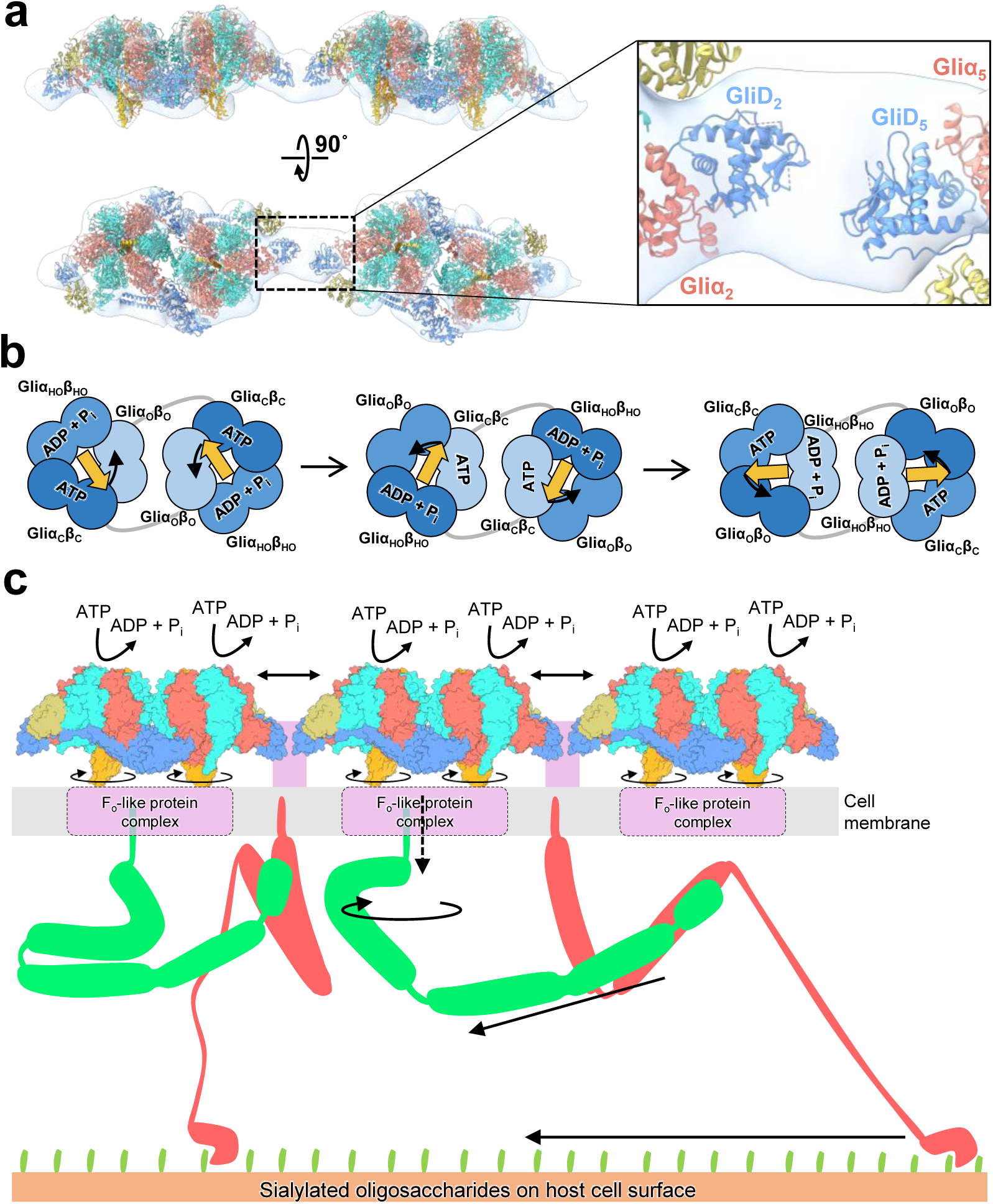
Structure and motion of the gliding machinery. (**a**) Fitting of the atomic model of the twin-motor into the EM map of the motor chain. The right panel indicates the interface between GliDs. (**b**) Possible rotational catalytic mechanism of F_1_-like ATPase. Gliγ is indicated by yellow arrows, the direction of which indicates the conformational tilt. (**c**) A possible explanation for the conversion mechanism from rotational to linear motion. Dotted arrows indicate unknown force transduction in the cell membrane. Gli349 and Gli521 are depicted in red and green, respectively. Unidentified protein regions in the gliding machinery are coloured purple.

## Discussion

The major difference between the twin motor and the dimeric form of eukaryotic mitochondrial ATP synthases^5^ is the interaction between the F_1_ domains through GliD and the N-terminal region of Gliβ. Although the GliD dimer does not have a transmembrane segment, the two long α-helices at the N-terminus are reminiscent of the structure of a peripheral stalk. This observation is consistent with the hypothesis that the MMOB1620 gene encoding GliD is derived through gene fusion from the b and δ subunits, which comprise the peripheral stalk^15^. The GliD-mediated dimer and chain formation may allow F_1_-like ATPase units to work cooperatively. Consistently, changes in the distance between twin motors have been associated with the ATP hydrolysis reaction, suggesting force transfer between the twin motors through GliD^3,4^. The PGK in the twin motor may efficiently supply ATP to the ATPase as fuel. Further, PGK molecules protruding from the transverse axis of the twin motor may be involved in the sheet formation of the internal chain, according to negative staining EM^25^. The weak density of PGK in the cryo-EM map can be explained by its flexibility, which was identified using high-speed atomic force microscopy^2^. The flexibility of the PGK molecule may suggest that the chain sheet serves as a cytoskeleton to withstand mechanical stress in the mycoplasma cell, which does not have a cell wall. The GliE subunit, which is bound to only one side of the dimeric F_1_-like ATPase, creates asymmetry in the twin motor structure and may contribute to the unidirectional nature of the gliding motility. However, this protein may also bind to another side of the dimeric F_1_-like ATPase in the cell but dislodge during purification.

The three distinct forms of the catalytic subunit Gliβ and their corresponding nucleotide binding suggest a rotary catalytic mechanism analogous to that of the rotary ATPases (Fig. 4b). In this rotary catalytic mechanism, Gliα_O_β_O_ transitions to Gliα_C_β_C_, and the DELSEED loop of Gliβ in this transition approaches Gliγ upon ATP binding at the catalytic site. The other Gliαβ dimers in the hexameric ring also operate cooperatively, with Gliα_C_β_C_ transitioning to Gliα_HO_β_HO_ upon ATP hydrolysis and Gliα_HO_β_HO_ transitioning to Gliα_O_β_O_ upon ADP and phosphate release. This one-step conformational change of the hexameric ring prompts a 120° rotation of Gliγ. Each Gliαβ dimer sequentially hydrolyses ATP, and through a three-step process, the hexameric ring structure and position of Gliγ revert to their initial states. The pseudo-two-fold rotational symmetry structure of the twin motor suggests that the two F_1_-like ATPase units operate in a synchronised manner, maintaining the structural symmetry. Throughout the catalytic reaction, significant changes in the positioning of the two F_1_-like ATPase units within the twin motor may be hindered due to the presence of the N-terminal regions of Gliβ bridging the two units and a buttress-like structure composed of two long α-helices of the GliD dimer (Fig. 2a, c, f).

For cell gliding, the rotary motions of the twin motors must be converted into a linear motion. This conversion mechanism may occur due to the membrane protein Gli521, which consists of a hook and rod, thereby functioning as a crank^26^ (Fig. 4c). Upon the conversion of the rotary motion to linear motion, Gli521 pulls the leg protein, Gli349, which is bound to sialylated oligosaccharides on the host cell surface, thereby propelling the mycoplasma cell forward^27^. The presence of a globular domain in Gliγ, similar to that in the F_1_-ATPase γ subunit, suggests a potential interaction with a membrane-embedded F_o_-like domain. MMOB1610 and MMOB1650, encoded in the Type 2 ATPase gene cluster, possess 12 and 2 transmembrane segments, respectively^15^. These two proteins are potential interaction partners for the globular domain of Gliγ at the cell membrane, transmitting the rotary motion to Gli521 on the cell surface.

The roles of rotary ATPases and their relatives, such as the flagellar type III protein export apparatus, which exhibit a hexameric ring with a central shaft, involve protein and ion transport^28,29^. We hypothesised that F_1_F_o_-ATPase has acquired the role and mechanism of the twin motor through an incidental contact with an adhesin on the membrane during the evolution of mycoplasma from a non-motile to a motile state^2,30^. This primitive gliding motility would have conferred a survival advantage. Another F_1_-like ATPase, referred to as Type 3 ATPase, is also found in mycoplasma, and this F_1_-like ATPase may drive the system that cleaves host antibodies^12,31^. Notably, Type 2 and 3 ATPases transition from F_1_-ATPase to drive mycoplasma-specific systems. Understanding these unique F_1_-like ATPases will contribute to investigations into the working principles and evolution of rotary ATPases.

## Methods

### Optical microscopy

The cells of the *M. mobile* mutant strain (gli521 [P476R]), which can glide like the wild type strain but binds sialylated oligosaccharides more tightly, were cultured, observed using phase-contrast microscopy, and recorded as previously described^32–35^. The recorded video was analysed using ImageJ software v1.54d (https://imagej.nih.gov/ij/).

### Isolation of the twin motor

The twin motor was isolated from the *M. mobile* mutant strain (gli521 [P476R]), as previously described^2^. The twin motor sample with phosphate-buffered saline (PBS) consisting of 8.1 mM Na_2_HPO_4_, 1.5 mM KH_2_PO_4_ (pH 7.3), 2.7 mM KCl and 137 mM NaCl, and 1 mM MgCl_2_ was concentrated to 1 mg/mL using a 100K Vivaspin concentrator (Merck, Germany) with 0.1% (w/v) CHAPS to prevent membrane adsorption. The concentrated twin-motor was subjected to 12.5% SDS-PAGE and stained using Coomassie brilliant blue R-250. The bands of the component proteins were identified using peptide mass fingerprinting, as previously described^36^.

### Cryo-EM grid preparation and data acquisition

An EG-grid was prepared as previously described^37^. A 2.6 μL sample solution was applied on the grid, automatically blotted from both sides with filter paper at 4 °C for 2 s at 100% humidity, and vitrificated using the semi-automated vitrification device, Vitrobot Mark IV (Thermo Fisher Scientific, USA). Cryo-EM imaging was performed using a CRYO ARM 300 (JEOL, Japan) operated at 300 kV with a K3 direct electron detector camera (AMETEK, Gatan, USA). A total of 7,350 movies were recorded using SerialEM software^38^ with a total dose of approximately 80 electrons Å^D2^ for 40 frames, an exposure time of 3.3 s per movie, and a nominal defocus range of 0.8 to 1.8 µm. The nominal magnification was 60,000×, corresponding to 0.87 Å per pixel.

### Cryo-EM image processing

The image processing steps for SPA are summarised in Extended Data Figs. 3 and 4. Image processing was performed using cryoSPARC v3.3.2^39^, unless otherwise stated. Movies were aligned using patch-based motion correction, and the contrast transfer function (CTF) parameters were estimated using patch CTF estimation. Micrographs with a CTF fit resolution worse than 8 Å were removed. Particles were automatically picked up using a Blob Picker, extracted with ×4 binning, and 2D-classified. The obtained 2D averaged images were used as templates for the Template Picker to pick up good particles. After two rounds of 2D classification, the selected good particles were subjected to local motion correction and re-extracted with the original pixel size. The particles were subjected to Ab-initio reconstruction using C1 symmetry enforced with a final resolution limit of 12 Å and heterogeneous reconstruction with 3 classes.

Particles in class 2 were subjected to homogeneous reconstruction using C1 symmetry, resulting in a map at a resolution of 3.5 Å. The final map was generated via local and global CTF refinement, and then non-uniform refinement. To improve the local resolution, the final map and particles were subjected to particle subtraction and local refinement using C1 symmetry to generate half-divided maps, each containing a monomeric ATPase. The whole and two local maps were subjected to mask creation and postprocessing in RELION 4.0^40^, resulting in resolutions of 3.2, 3.1, and 3.1 Å, respectively. Local resolution estimation was also performed in RELION 4.0. The post-processed map was denoised using Topaz software v0.2.5 with the trained model Unet-3d-10a^41^. RELION 4.0 was used to calculate the Fourier shell correlation (FSC) curve between two half-maps and estimate the local resolution.

### Modelling

Initial models for GliαβγDE were generated using AlphaFold2^42^, whereas the homology model for PGK was generated using Modeller v10.4^43^ with the atomic model of PGK (PDB ID: 4DG5) from *S. aureus*. The generated models were fitted to the corresponding densities of the EM map as rigid bodies using UCSF Chimera v1.15^44^. The fitted models were manually inspected and adjusted using COOT v0.9.8.1^45^ and refined using the *phenix.real_space_refine* program in PHENIX v1.19^46,47^ with reference model restraints using atomic models of F_1_-ATPase (PDB ID: 5IK2) from *Caldalaklibacillus thermarum* and PGK (PDB ID: 3ZLB) from *Streptococcus pneumoniae*. Manual adjustment using COOT and refinement using PHENIX were repeated until model parameters were no longer improved. FSC curves between the map and model were calculated using PHENIX. The refined model was also evaluated via comprehensive validation in PHENIX and the Q-score^48^. To identify the nucleotide binding sites of F_1_-like ATPases, F_o_-F_c_ maps were generated using Servalcat^49^. Superposition of models and fitting of models to the map were performed using UCSF ChimeraX v1.2.5^50^. All structural figures were obtained using UCSF ChimeraX.

## Data availability

The cryo-EM map for the twin motor is available on the Electron Microscopy Data Bank (EMDB) under accession code: EMD-60718. The coordinates have been deposited in the Protein Data Bank (PDB) under accession code: 9IO5.

### Acknowledgments

We thank Tomomi Shimonaka at Osaka Metropolitan University and Fumiaki Makino at Osaka University for their technical assistance. This manuscript was edited using Copilot, an AI tool. This study was supported by Grants-in-Aid for Scientific Research (A) (JP17H01544), a JST CREST grant (JPMJCR19S5) to M.M., and Research Support Project for Life Science and Drug Discovery (BINDS) from AMED under Grant Number, JP22am121003, to K.N., and JEOL YOKOGUSHI Research Alliance Laboratories of Osaka University to K.N.

## Author contributions

T.T. and M.M. conceived the strategy. M.M. and K.N. supervised the research. T.T. prepared the cryo-EM samples and carried out the biochemical analyses. J.F. prepared EG-grid for cryo-EM observation. A.K. and T.M. performed cryo-EM. T.T. carried out 3D reconstruction and model building. T.K., T.H., A.K., T.M., K.K., and K.N. participated in the discussion regarding the structural analyses. T.T. and M.M. wrote the manuscript with input from all authors.

