## Extended Data Fig. 1-15, Table 1-3 for "Dimeric assembly of F_1_-like ATPase for the gliding motility of *Mycoplasma*"

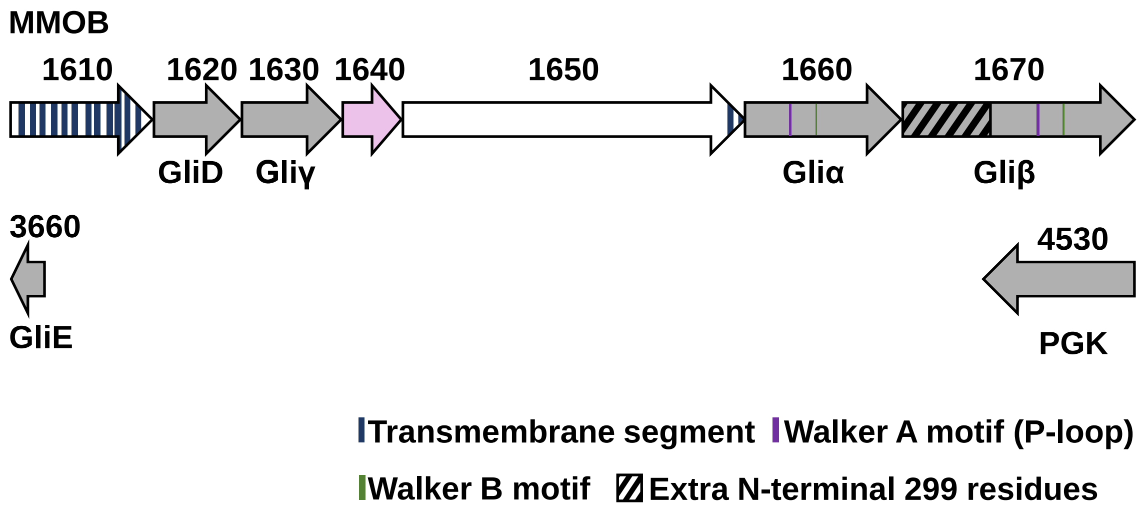


**Extended Data Fig. 1 Open reading frames related to twin motor in *M. mobile*.**

The Type 2 ATPase cluster is composed of MMOBs 1610–1670. The twin-motor components are shown in gray. In this study, MMOB1640, coloured in pink, was suggested as a component only by SDS-PAGE analysis.

a

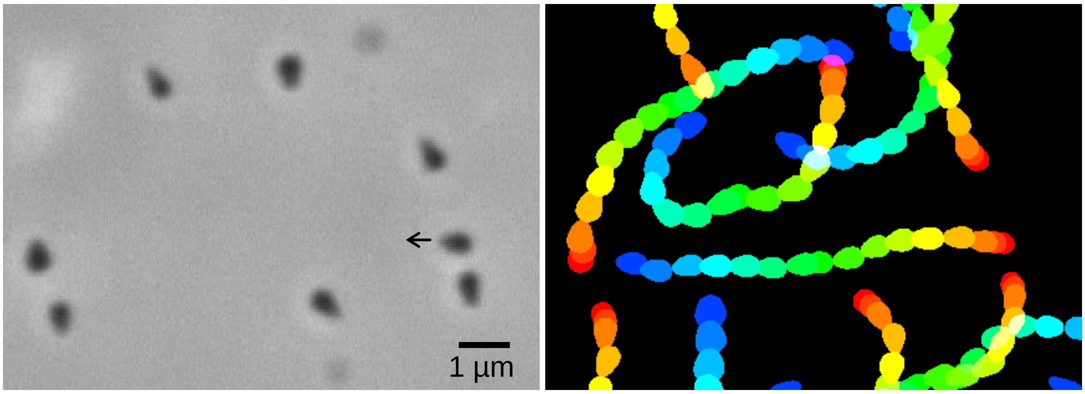


**Extended Data Fig. 2 Optical microscopy of *M. mobile* gliding.**

The left panel shows the cell image. Arrow indicates the gliding direction. The right panel shows the rainbow trace of *M. mobile* gliding. The gliding cells were traced every 0.2 s and stacked for 3 s. The trajectories are coloured from red to blue.

| | Twin motor (EMD-60718) (PDB ID: 9IO5) |
|---|---|
| **Data collection and processing** | |
| Microscope | CRYO ARM 300 |
| Detector | K3 |
| Nominal magnification | 60,000× |
| Voltage (kV) | 300 |
| Total electron exposure ($e^-$/Å$^2$) | 80 |
| Frames (no.) | 40 |
| Total exposure time (sec) | 3.3 |
| Defocus range (µm) | −0.8 to −1.8 |
| Pixel size (Å) | 0.87 |
| Total image sets (no.) | 7,350 |
| Used image sets (no.) | 7,074 |
| Initial particle images (no.) | 633,820 |
| Final particle images (no.) | 142,490 |
| Symmetry imposed | C1 |
| Map resolution (Å) | 3.2 |
| B-factor for sharpening (Å$^2$) | −57.3 |
| FSC threshold | 0.143 |
| **Model refinement** | |
| Initial models used | Homology models and AlphaFold2-predicted structures |
| Model resolution | 3.2 |
| FSC threshold | 0.5 |
| Model composition | |
| Non-hydrogen atoms (no.) | 69,678 |
| Protein residues (no.) | 8,930 |
| Ligands (no.) | 22 |
| B factors | |
| Protein (Å$^2$) | 35.37 |
| Ligands (Å$^2$) | 34.50 |
| R.m.s.deviations | |
| Bond lengths (Å$^2$) | 0.004 |
| Bond angles (˚) | 0.944 |
| Valiation | |
| Molprobity score | 1.48 |
| Clashscore | 5.46 |
| Poor rotamers (%) | 0.00 |
| Ramachandran plot | |
| Favored (%) | 96.91 |
| Allowed (%) | 3.06 |
| Disallowed (%) | 0.03 |

**Extended Data Table 1 Cryo-EM data collection and model statistics.**

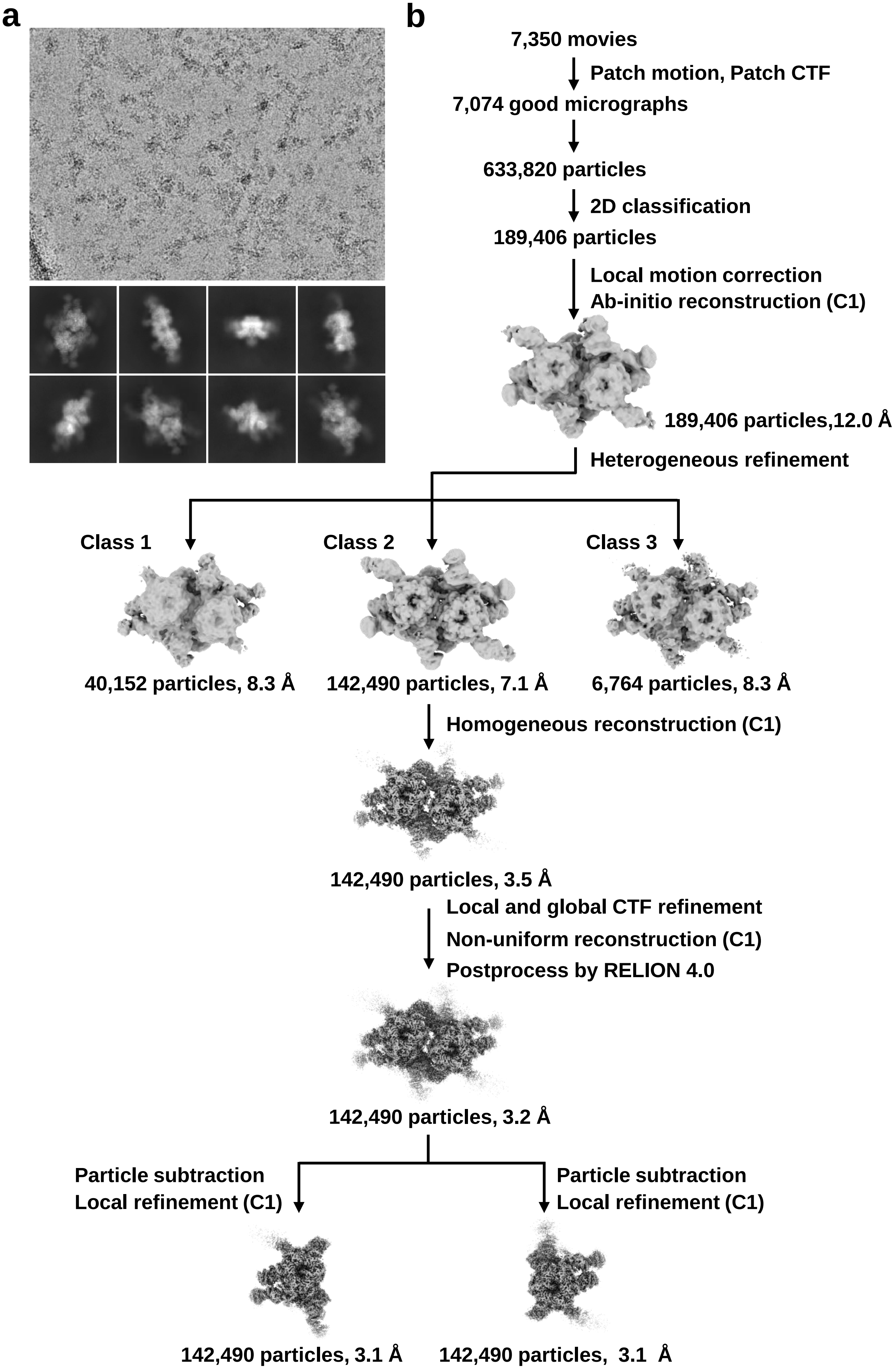
a
b
7,350 movies
Patch motion, Patch CTF
7,074 good micrographs
633,820 particles
2D classification
189,406 particles
Local motion correction
Ab-initio reconstruction (C1)
189,406 particles,12.0 Å
Heterogeneous refinement
Class 1
Class 2
Class 3
40,152 particles, 8.3 Å
142,490 particles, 7.1 Å
6,764 particles, 8.3 Å
Homogeneous reconstruction (C1)
142,490 particles, 3.5 Å
Local and global CTF refinement
Non-uniform reconstruction (C1)
Postprocess by RELION 4.0
142,490 particles, 3.2 Å
Particle subtraction
Local refinement (C1)
Particle subtraction
Local refinement (C1)
142,490 particles, 3.1 Å
142,490 particles, 3.1 Å

**Extended Data Fig. 3 Image processing.**

(a) Representative micrograph (top) and averaged images (bottom). (b) Workflow of image processing.

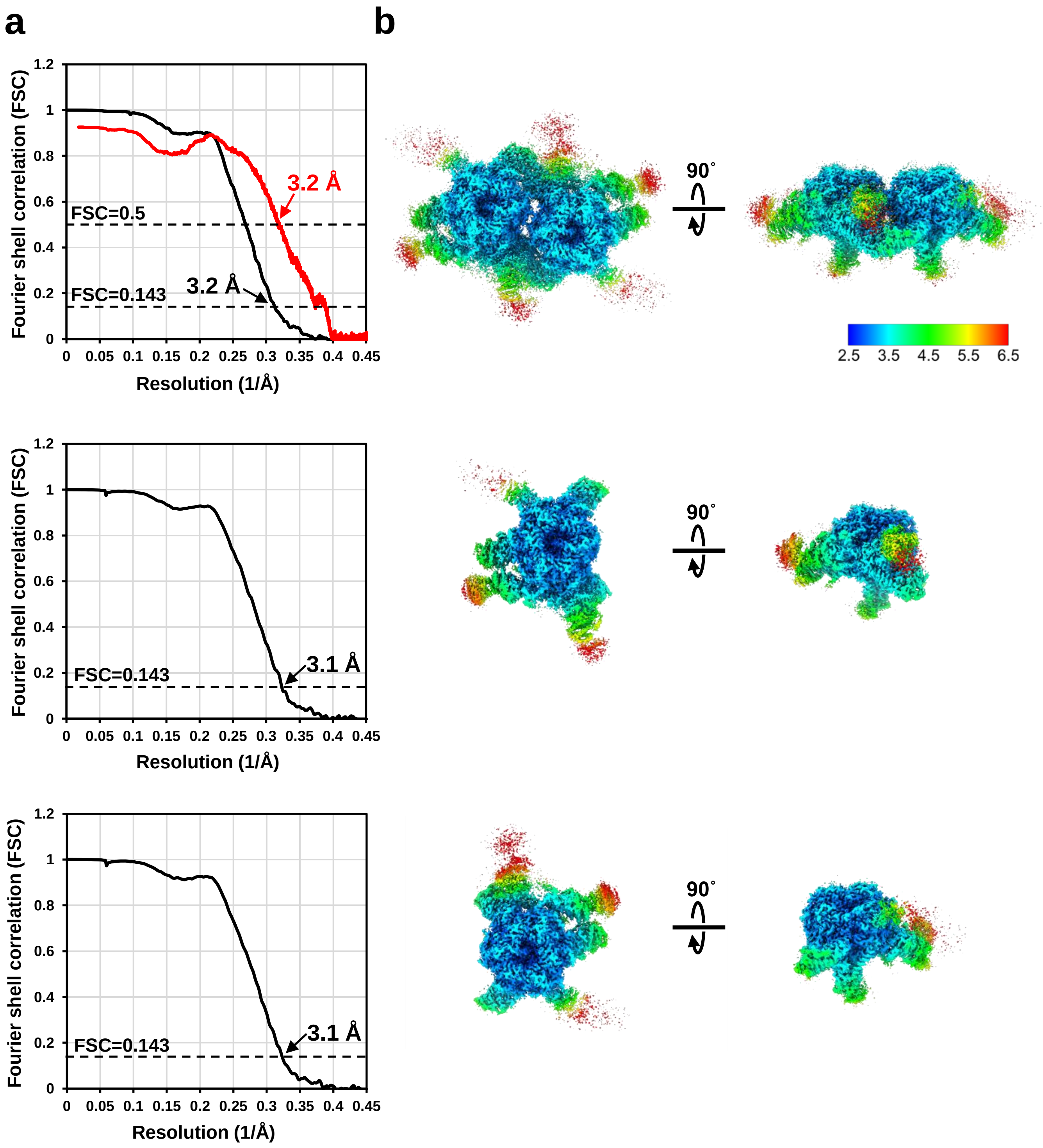


**Extended Data Fig. 4 Fourier shell correlation and local resolution estimation of density maps.**

(a) Fourier shell correlation curves for half maps (black) and cross-validation between the density map and model (red). (b) Local resolution estimated using RELION.

| Chain ID | Protein | Protein length (amino acids) | Modeled amino acid residues | Q-score |
|---|---|---|---|---|
| A | Gliβ$_1$ | 784 | 281–778 | 0.60 |
| B | Gliβ$_2$ | 784 | 202–771 | 0.59 |
| C | Gliβ$_3$ | 784 | 217–777 | 0.57 |
| D | Gliα$_1$ | 528 | 1–528 | 0.61 |
| E | Gliα$_2$ | 528 | 1–528 | 0.58 |
| F | Gliα$_3$ | 528 | 1–528 | 0.60 |
| G | Gliγ$_1$ | 336 | 1–56, 76–190, 243–331 | 0.49 |
| H | Gliβ$_4$ | 784 | 281–779 | 0.60 |
| I | Gliβ$_5$ | 784 | 201–771 | 0.57 |
| J | Gliβ$_6$ | 784 | 125–159, 213–774 | 0.58 |
| K | Gliα$_4$ | 528 | 1–528 | 0.60 |
| L | Gliα$_5$ | 528 | 1–528 | 0.58 |
| M | Gliα$_6$ | 528 | 1–528 | 0.60 |
| N | Gliγ$_2$ | 336 | 1–57, 77–190, 247–331 | 0.48 |
| O | PGK$_1$ | 511 | 179–344, 348–385 | 0.37 |
| P | PGK$_2$ | 511 | 179–365, 370–385 | 0.34 |
| Q | PGK$_3$ | 511 | 179–343, 349–386 | 0.33 |
| R | GliD$_1$ | 293 | 47–53, 58–270 | 0.47 |
| S | GliD$_2$ | 293 | 109–194, 207–249, 255–272, 277–287 | 0.36 |
| T | GliD$_3$ | 293 | 62–290 | 0.55 |
| U | GliD$_4$ | 293 | 45–270 | 0.50 |
| V | GliD$_5$ | 293 | 106–195, 207–288 | 0.38 |
| W | GliD$_6$ | 293 | 65–290 | 0.52 |
| X | GliE | 112 | 4–112 | 0.59 |
| Y | Unknown helix | | 1–13 | 0.45 |
| Z | Unknown helix | | 1–9 | 0.35 |

**Extended Data Table 2 Calculated Q-score for each subunit.**

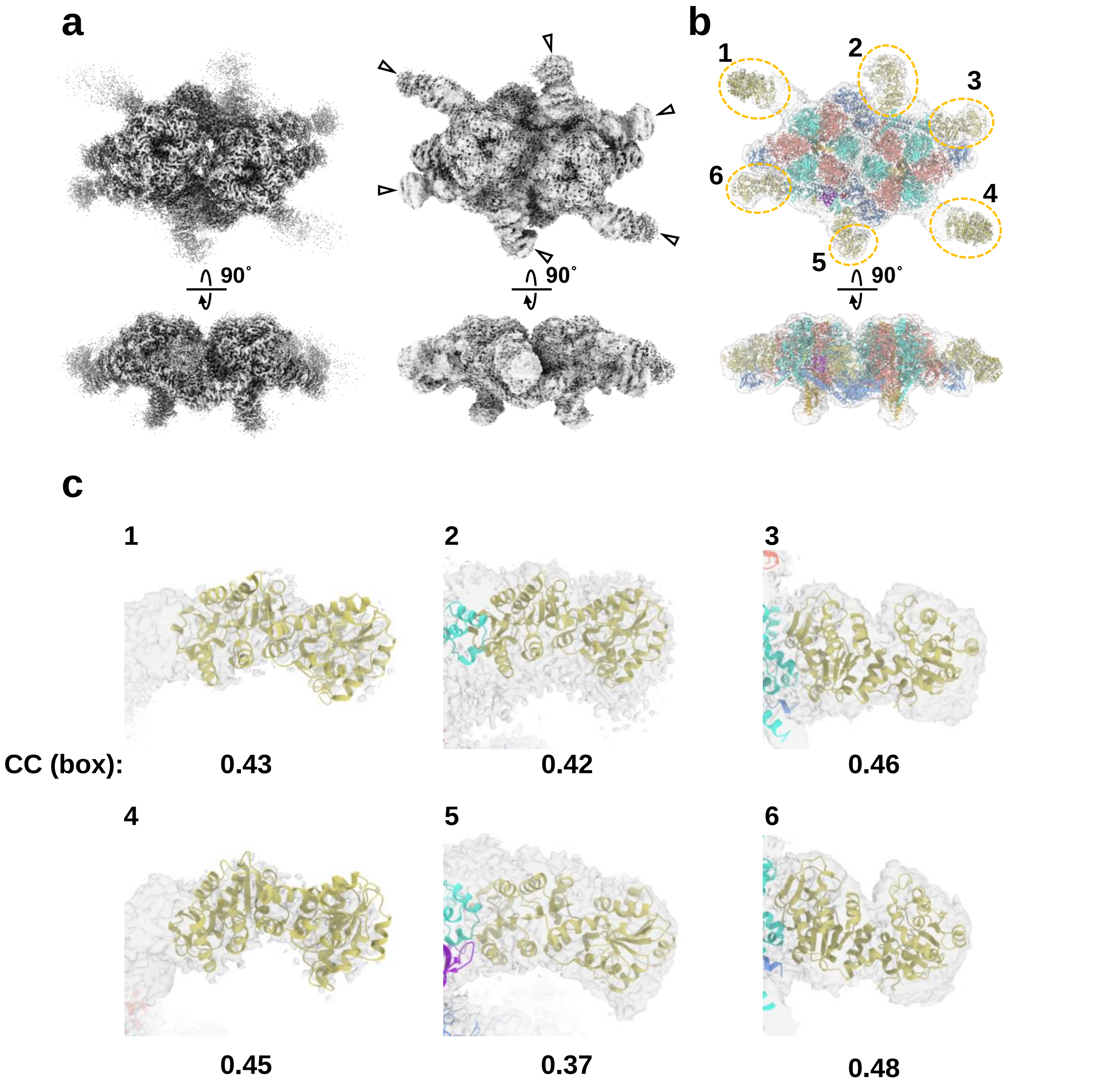


**Extended Data Fig. 5 Densities corresponding to PGK molecules.**

(a) Comparison between maps before and after denoising (left and right). Densities due to denoising are indicated by triangles. The maps before and after denoising are contoured at 0.55 and 0.12 in UCSF ChimeraX. (b) Denoised map fitted with atomic models of twin-motor subunits and PGKs from *S. aureus* (PDB ID: 4DG5). Densities corresponding to the PGK molecules were marked with orange dot circles. (c) Fitting of PGK molecules; numbers correspond to (b).

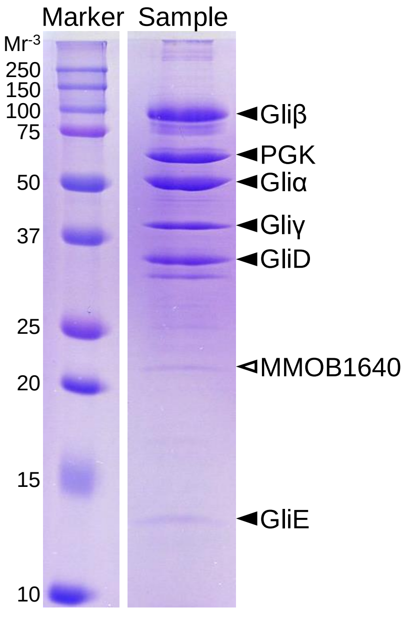


**Extended Data Fig. 6 SDS-PAGE of isolated twin motor.**

The sample was subjected to 12.5% SDS-PAGE gel and stained with Coomassie brilliant blue R-250. Twin-motor components are indicated by black triangles. The possible component is indicated by a white triangle. Molecular masses are shown on the left.

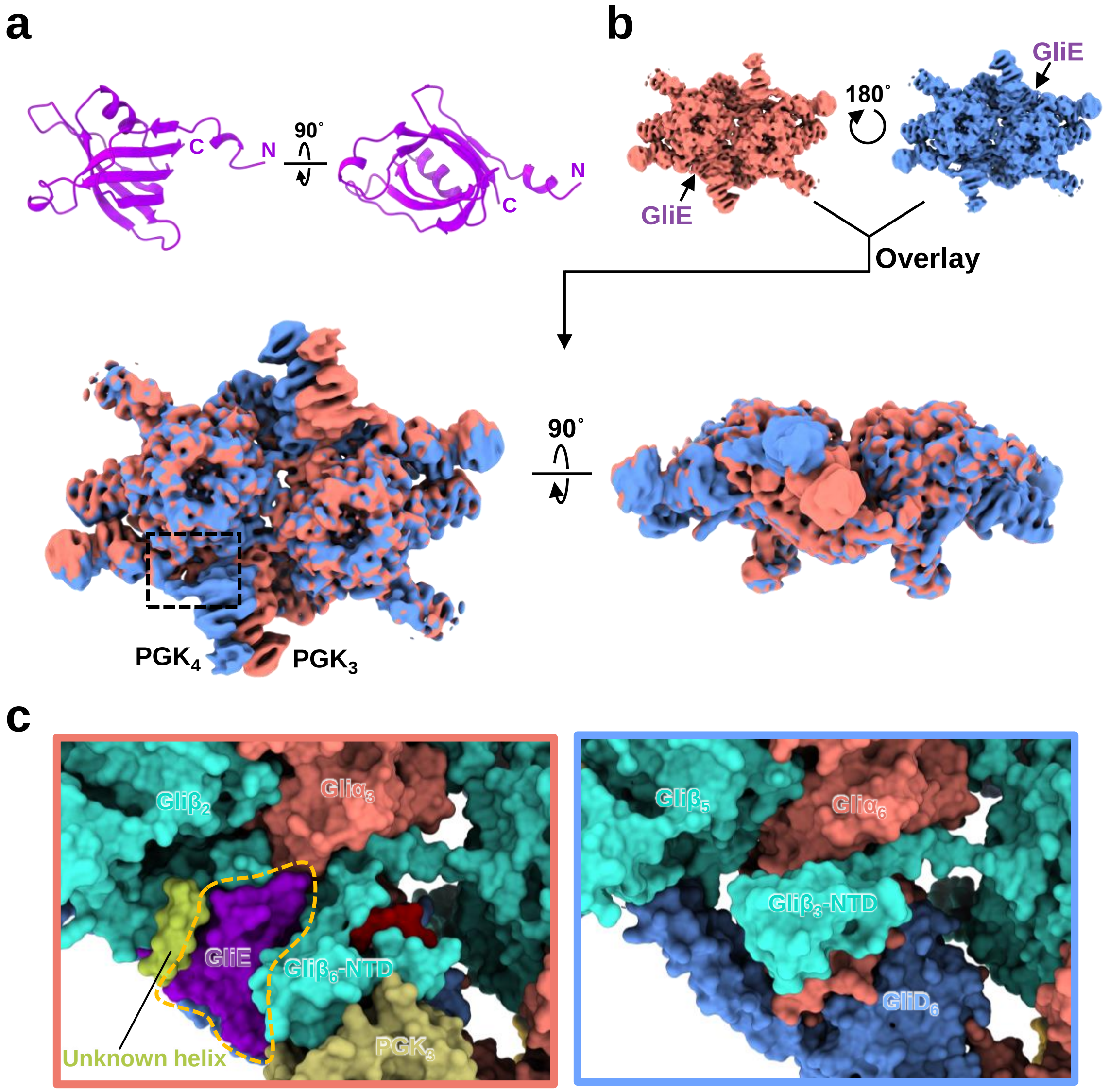


**Extended Data Fig. 7 Asymmetry of twin-motor structure.**

(a) GliE structure. (b) Overlay of the twin-motor structures rotated by 180˚. The maps were low-pass filtered at 8 Å in RELION and contoured at 0.17 in UCSF ChimeraX. (c) Comparison of rotated twin-motor structures. The dotted square area in (b) is shown. GliE is surrounded by an orange dot line.

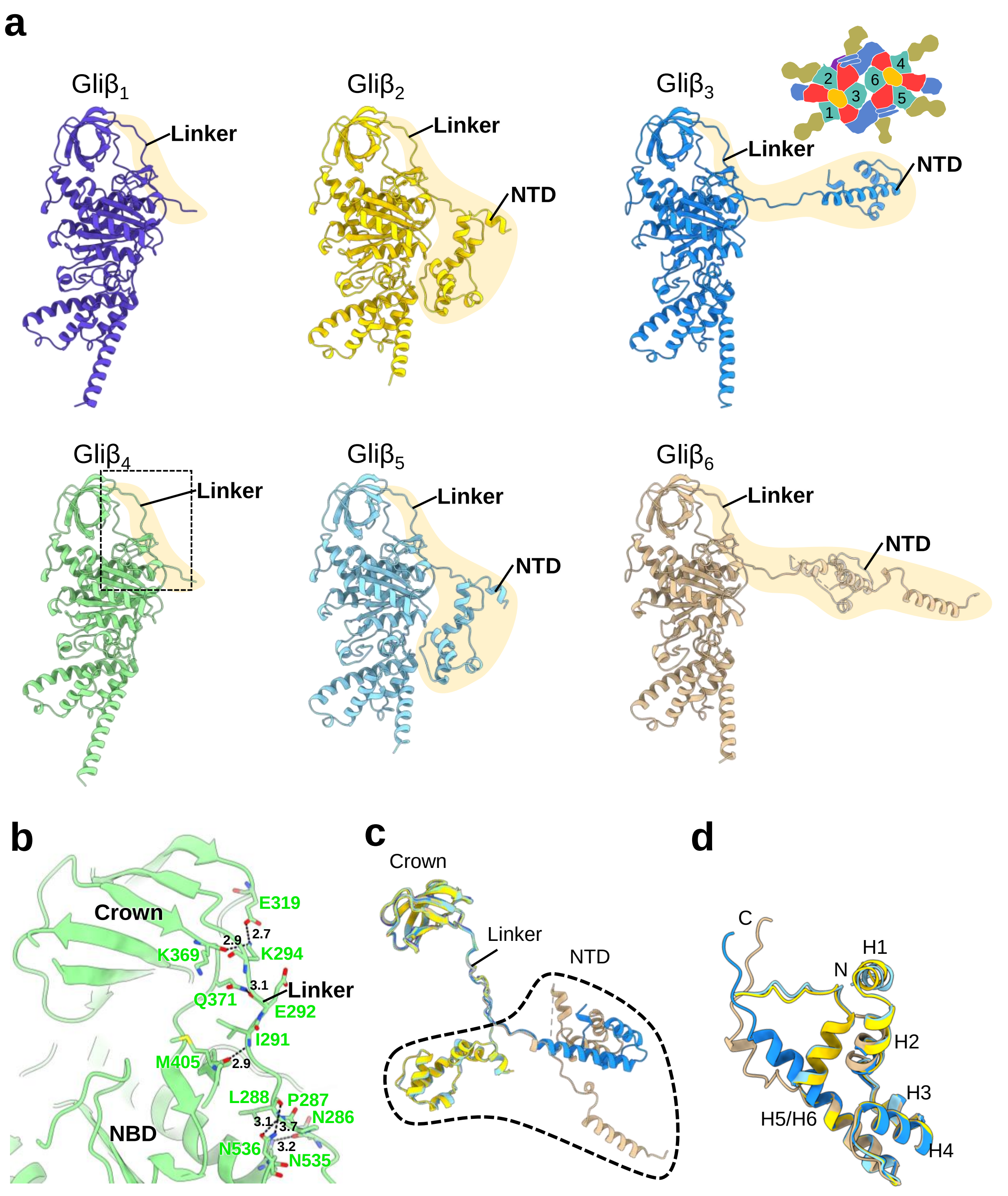


**Extended Data Fig. 8 Extended N-terminal region of Gliβ.**

(a) Structures of Gliβ$_{1\text{-}6}$ in twin motor. Extended N-terminal regions are marked in khaki. An illustration with the location of each Gliβ is presented in the upper right corner. (b) Linker interactions with crown and NBD in Gliβ$_4$.The region corresponds to the black dot box in (a). (c) Superposition of structures composed of crown, linker, and NTD. Structures were superimposed on the crown. (d) Superposition of NTD.

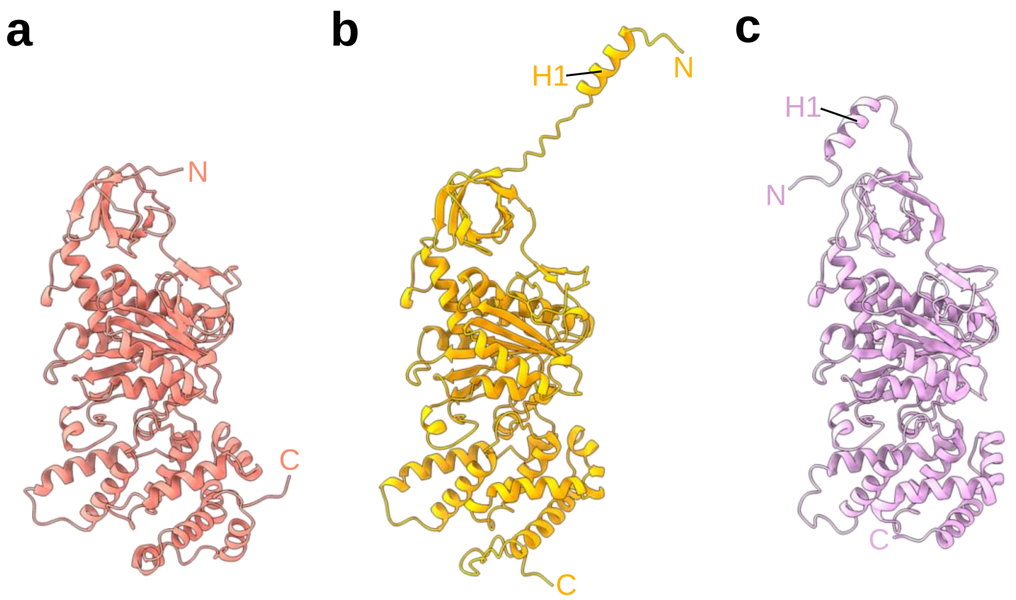


**Extended Data Fig. 9 N-terminal helix (H1) of $F_1$-ATPase α not found in Gliα.** (a) Gli$\alpha_2$ structure. (b) Structure of $F_1$-ATPase (Type 1 ATPase) α from *M. mobile* predicted using AlphaFold2. (c) Structure of $F_1$-ATPase α from *Bacillus* PS3 (PDB ID: 6N2Y).

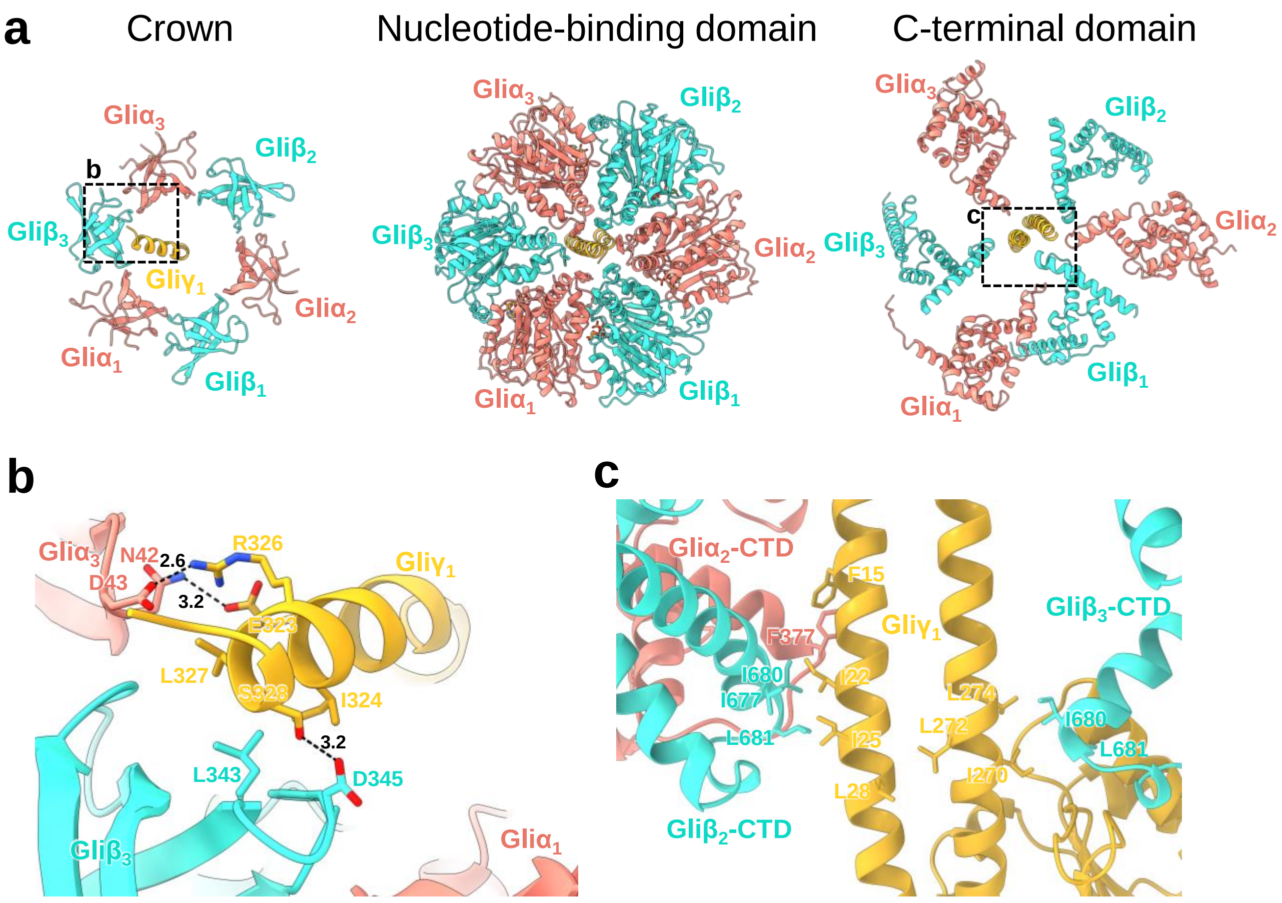


**Extended Data Fig. 10 Asymmetry of $F_1$-like ATPase and interaction between hexameric ring and Gliγ.**

(a) Cross-section of each domain of $F_1$-like ATPase. (b) Interaction between the crown region of the hexameric ring and extended helix of Gliγ. The region corresponds to the black dot box in (a). (c) Hydrophobic interaction between the C-terminal domain of the hexameric ring and coiled-coil of Gliγ. The region corresponds to the black dot box in (a).

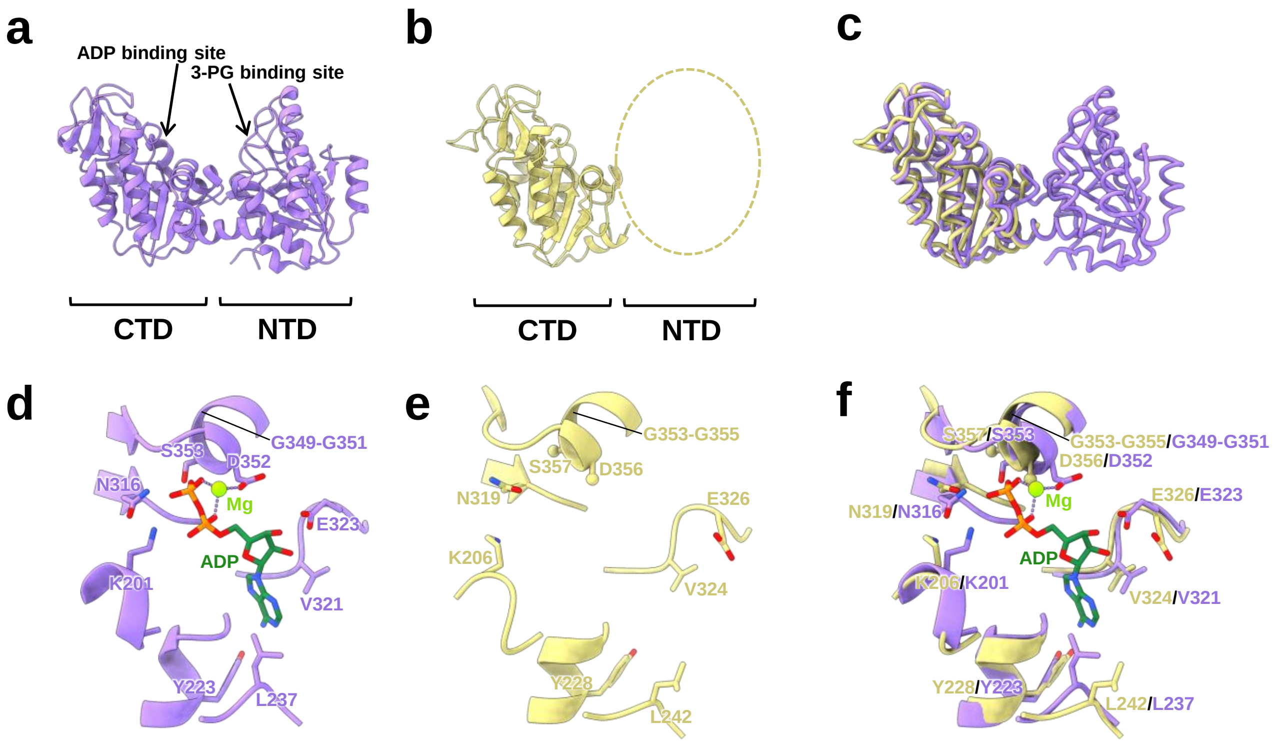


**Extended Data Fig. 11 Comparison of the PGK structure.**

(a) PGK structure from *S. aureus* (PDB ID: 4DG5). (b) PGK structure in the twin motor. (c) Superimposition of the PGK molecules. (d) Catalytic site binding Mg-ADP in PGK (PDB ID: 1PHP). (e) Catalytic site of PGK in the twin motor. (f) Superimposition of the catalytic sites.

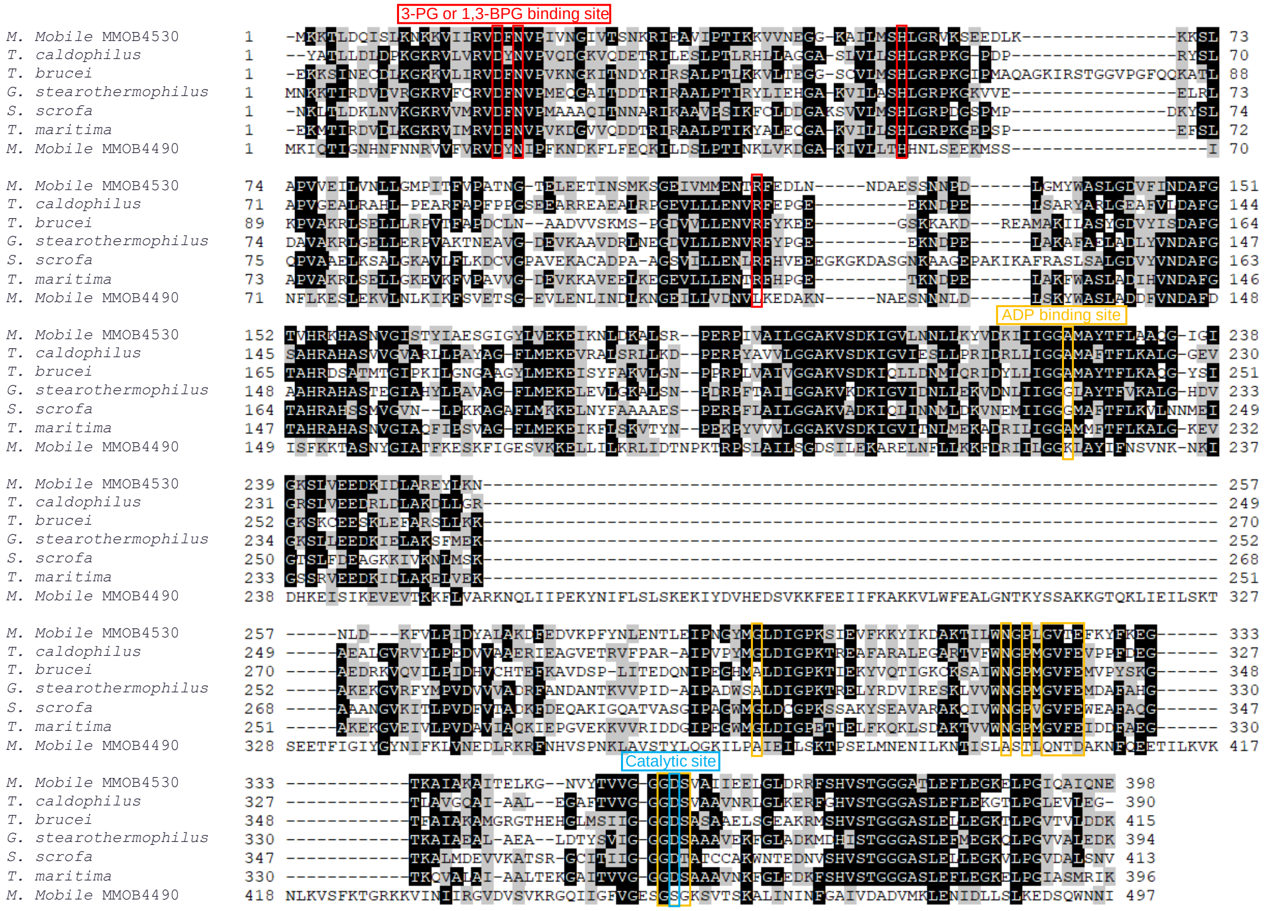

**Extended Data Fig. 12 Amino acid sequence alignment of PGK and the homologue.**

ADP binding, catalytic, and 3-PG or 1,3-BPG binding sites are indicated by orange, blue, and red boxes, respectively. Amino acid sequences of PGKs and the homologue were aligned using ClustalW. The figure was generated using BioEdit.

| | 0° (*ATP-waiting*) | 0° (*step-waiting*) | 81° | 83° (*post-hyd*) | 91° | 101° |
|---|---|---|---|---|---|---|
| Gli$\beta_1$ (C) | 2.05 | 2.16 | 2.18 | 2.15 | 2.15 | 2.16 |
| Gli$\beta_2$ (HO) | 1.95 | 2.26 | 3.36 | 2.67 | 2.67 | 2.03 |
| Gli$\beta_3$ (O) | 2.19 | 2.18 | 2.33 | 2.33 | 2.33 | 2.35 |

**Extended Data Table 3 RMSD values (Å) between Gliβ and *Bacillus* PS3 $F_1$-ATPase β.**

Gli$\beta_{1\text{-}3}$ (C, HO, and O) were superimposed on the same conformation of the β subunit in the six states of *Bacillus* PS3 $F_1$-ATPase (PDB ID: 8HH1–8HH6), respectively. The state names of *Bacillus* PS3 $F_1$-ATPase are based on the γ subunit position[8]. The lowest RMSD value for each Gliβ is shown in red.

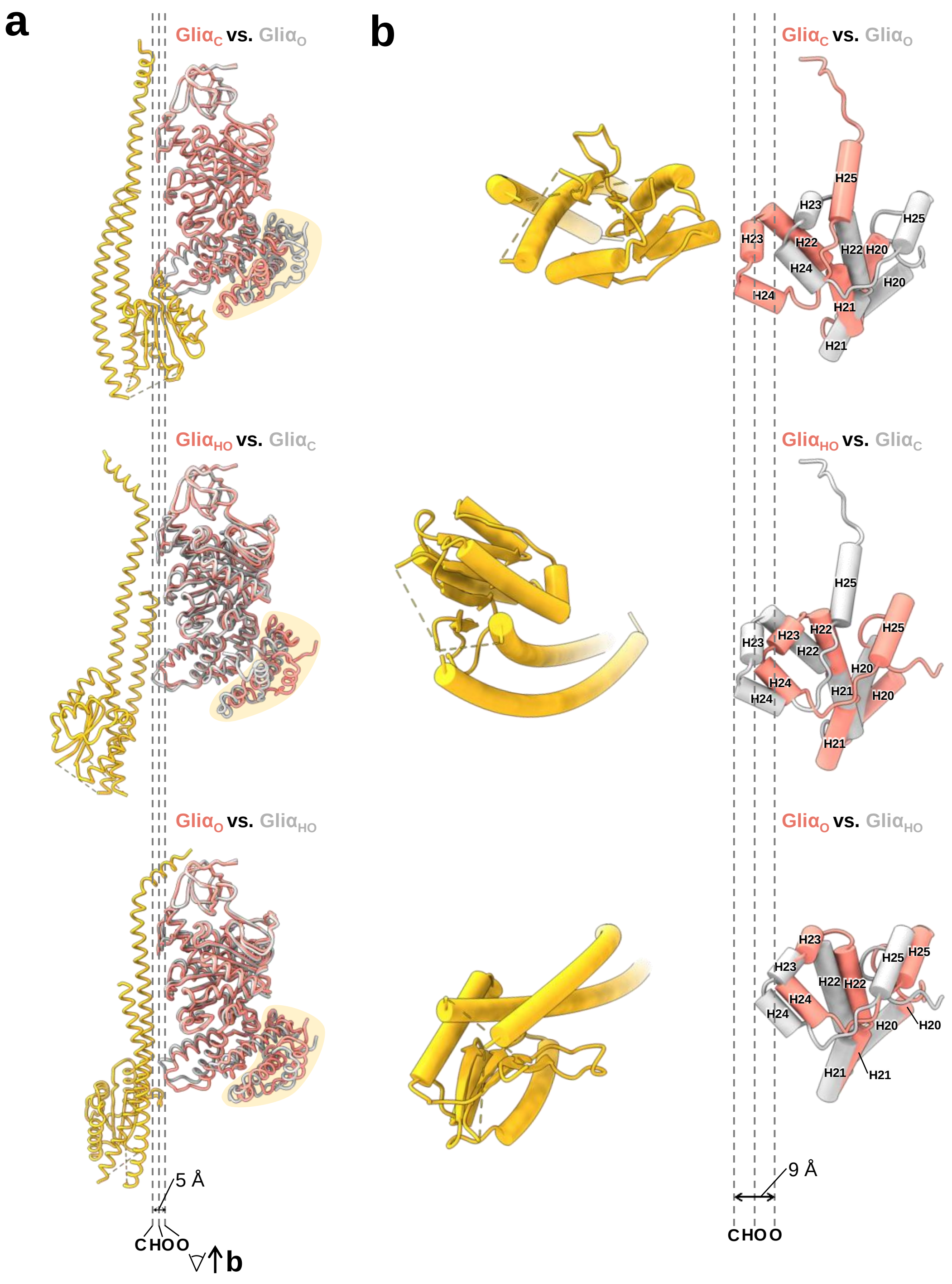


**Extended Data Fig. 13 Differences in the conformations of the three Gliα.**

(a) Superposition of three Gliα in the $F_1$-like ATPase. The dotted lines indicate the position of the region corresponding to DELSEED loop in each Gliα. H20–H25 regions are marked in khaki. (b) Comparison of the H20–H25 region among the three Gliα. Each image is a bottom-view of (a).

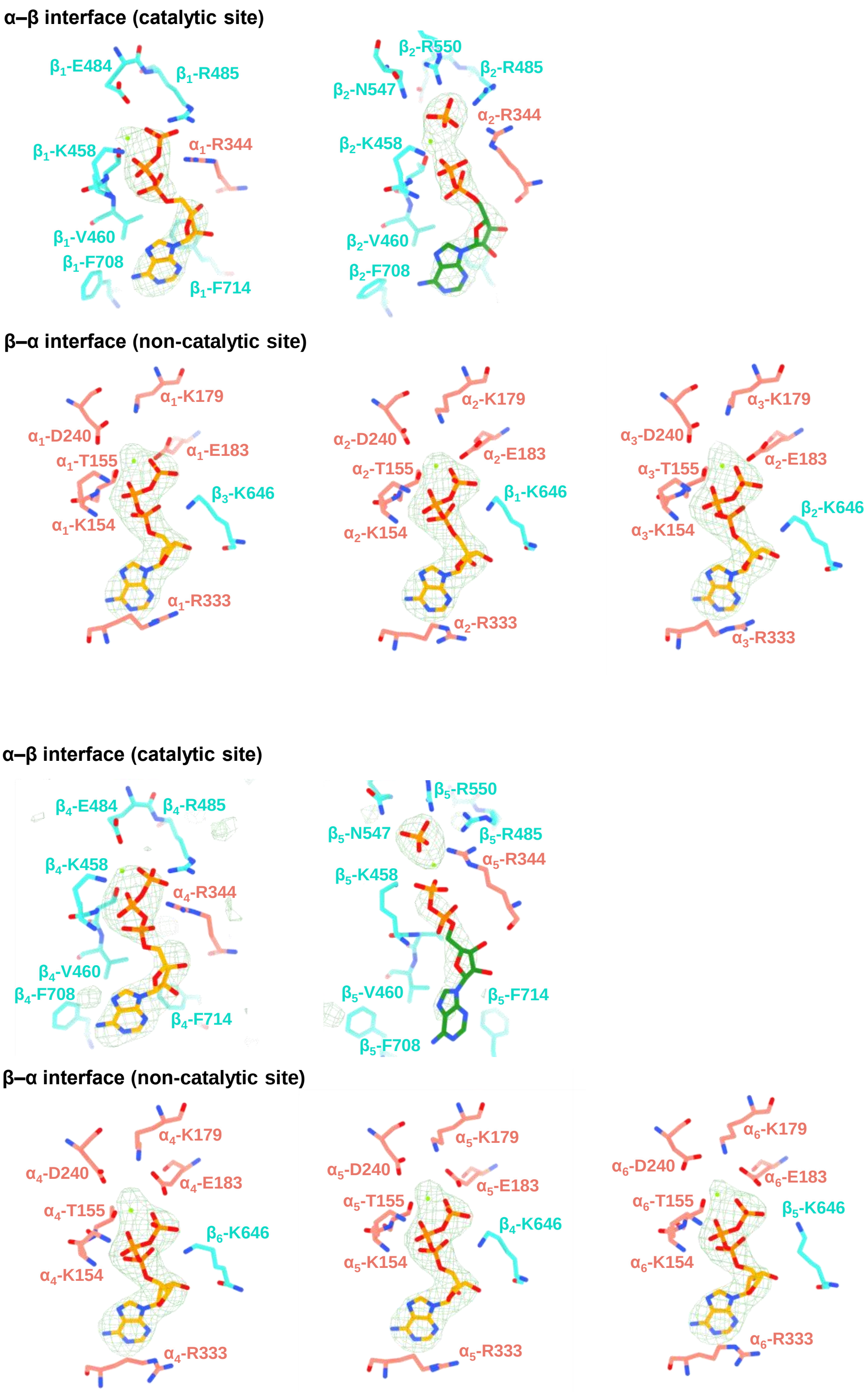


**Extended Data Fig. 14 $F_o$-$F_c$ maps of the nucleotide binding sites of $F_1$-like ATPases.**

The maps are contoured at 5.50 except for the Gli$\alpha_4\beta_4$ (3.30) and Gli$\alpha_5\beta_5$ (4.00) interfaces in UCSF ChimeraX.

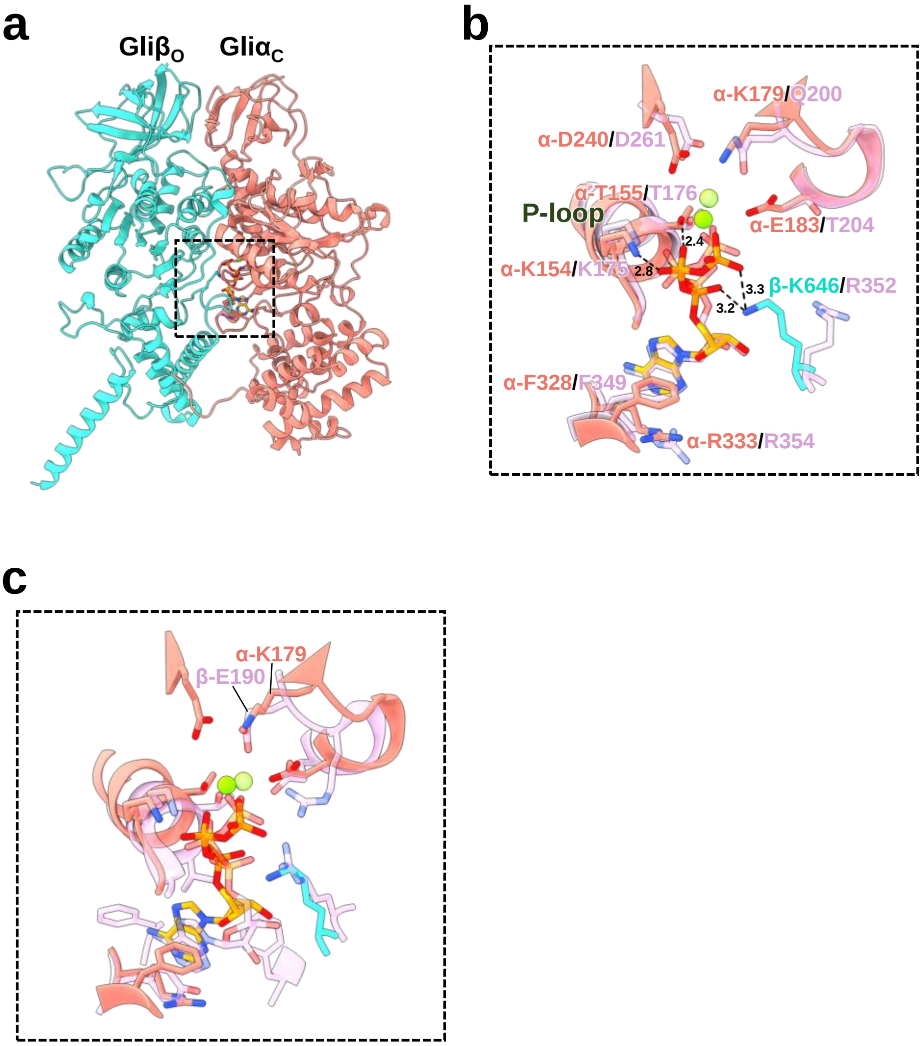


**Extended Data Fig. 15 Interface corresponding to the non-catalytic site of $F_1$-ATPase.**

(a) Interface formed by Gliβ$_O$ and Gliα$_C$. (b, c) Comparison of the interface between the $F_1$-ATPase and $F_1$-like ATPase. The interface from the dot square in (a) is superimposed on the corresponding non-catalytic site (b) and catalytic site (c) of $F_1$-ATPase (PDB ID: 8HHA), coloured purple.
